# Quality control strategies for brain MRI segmentation and parcellation: practical approaches and recommendations - insights from The Maastricht Study

**DOI:** 10.1101/2021.02.01.428681

**Authors:** Jennifer Monereo Sánchez, Joost J.A. de Jong, Gerhard S. Drenthen, Magdalena Beran, Walter H. Backes, Coen D.A. Stehouwer, Miranda T. Schram, David E.J. Linden, Jacobus. F.A. Jansen

## Abstract

**Background:** Quality control of brain segmentation is a fundamental step to ensure data quality. Manual quality control is the current gold standard, despite unfeasible in large neuroimaging samples. Several options for automated quality control have been proposed, providing potential time efficient and reproducible alternatives. However, those have never been compared side to side, which prevents to reach consensus in the appropriate QC strategy to use. This study aims to elucidate the changes manual editing of brain segmentations produce in morphological estimates, and to analyze and compare the effects of different quality control strategies in the reduction of the measurement error.

**Methods:** We used structural MR images from 259 participants of The Maastricht Study. Morphological estimates were automatically extracted using FreeSurfer 6.0. A subsample of the brain segmentations with inaccuracies was manually edited, and morphological estimates were compared before and after editing. In parallel, 11 quality control strategies were applied to the full sample. Those included: a manual strategy, manual-QC, in which images were visually inspected and manually edited; five automated strategies where outliers were excluded based on the tools MRIQC and Qoala-T, and the metrics morphological global measures, Euler numbers and Contrast-to-Noise ratio; and five semi-automated strategies, were the outliers detected through the mentioned tools and metrics were not excluded, but visually inspected and manually edited. We used a regression of morphological brain measures against age as a test case to compare the changes in relative unexplained variance that each quality control strategy produces, using the reduction of relative unexplained variance as a measure of increase in quality.

**Results:** Manually editing brain surfaces produced changes particularly high in subcortical brain volumes and moderate in cortical surface area, thickness and hippocampal volumes. The exclusion of outliers based on Euler numbers yielded a larger reduction of relative unexplained variance for measurements of cortical area, subcortical volumes and hippocampal subfields, while manual editing of brain segmentations performed best for cortical thickness. MRIQC produced a lower, but consistent for all types of measures, reduction in relative unexplained variance. Unexpectedly, the exclusion of outliers based on global morphological measures produced an increase of relative unexplained variance, potentially removing more morphological information than noise from the sample.

**Conclusion:** Overall, the automatic exclusion of outliers based on Euler numbers or MRIQC are reliable and time efficient quality control strategies that can be applied in large neuroimaging cohorts.

## 1 Introduction

Quality control (QC) of brain MRI segmentation and parcellation (i.e. the detection and correction or exclusion of inaccuracies in segmented brain images), is a fundamental step to ensure measurement reliability. The concept of QC has recently gained interest, with many tools, metrics, and protocols being proposed (Backhausen et al., 2016; Esteban et al., 2017; Keshavan et al., 2018; Klapwijk et al., 2019; Rosen et al., 2018; Waters et al., 2019). Manual QC, consisting of visual inspection and surface editing, despite its component of subjectivity, is considered the gold standard. However, neuroimaging research is shifting towards Big Data paradigms, with studies including thousands of brain images, such as the UK Biobank (Miller et al., 2016), or The Maastricht Study (Schram et al., 2014). Manual QC has therefore become an unfeasible strategy due to the time and resources required, thus there is a need to validate and come to an agreement in a reproducible and time efficient QC solution for large cohort studies.

Poor quality segmentation is defined by the presence of certain amount or severity of inaccuracies. Inaccuracies occur when the boundaries that define the morphological divisions or regions of interest (ROIs) do not correspond to the anatomical boundaries, which may lead to morphological measurement errors. A commonly used tool to segment structural brain MRI is FreeSurfer (Fischl, 2012), a software for MRI analysis that provides automated subcortical segmentation and cortical parcellation of the brain. Errors in FreeSurfer’s output may happen (amongst others) when sufficiently abnormal brain structure, or low quality image is provided as input. Further, image artifacts have been related to worse segmentation estimates for both cortical thickness (Reuter et al., 2015) and volumes (Savalia et al., 2017). Errors in the segmentation may result in regression attenuation (Hutcheon et al., 2010), as well as reduction of statistical power (Phillips and Jiang, 2016) in regression analysis of phenotypical measures with MR features. Large sample sizes can compensate for these downsides. However, when the measurement errors are systematic, recurrent, and in the same direction, a bias can be introduced, making segmentation quality a potential confounder. This type of bias has previously been shown in clinical populations compared to healthy controls (Pardoe et al., 2016), children compared to adults (Blumenthal et al., 2002) and older adults compared to younger adults (Madan, 2018; Savalia et al., 2017; Wenger et al., 2014).

Manual QC of brain segmentations is currently the most accepted approach to ensure reliable segmentation estimates, and in absence of a better solution, it is considered the gold standard for QC. The manual QC process involves the visual inspection of each segmentation, ideally by several independent operators with knowledge of neuroimaging normal anatomy, in addition to the manual editing of segmentations identified as inaccurate. This process is time consuming (the time required to visually inspect and edit each segmentation can range between 10 and 45 minutes) and subjective, requiring a trained operator. Moderate interrater reliability has been previously reported, with Cohen’s Kappa indices that range between 0.30 (Esteban et al., 2017) and 0.48 (Savalia et al., 2017).

Studies investigating alterations in morphological estimates due to manual editing have shown mixed results, with some studies reporting significant changes (Beelen et al., 2020; Waters et al., 2019), while others showed no differences (McCarthy et al., 2015). Despite the potential changes in morphological estimates due to manual editing, manual QC does not show important effects in the sensitivity to detect differences between groups using volumetric morphological estimates (Waters et al., 2019), although it may have more impact on the sensitivity to detect differences using cortical morphological estimates like surface area and thickness (Beelen et al., 2020).

Rather than performing manual QC, one can also use automatic exclusion of cases based on quantitative parameters, such as quality metrics or morphological information that are readily available from FreeSurfer or other software, making the QC process more time-efficient. Among the most commonly used quality metrics are contrast-to-noise ratio (CNR) (Welvaert and Rosseel, 2013) and Euler numbers (EN) (Dale et al., 1999). CNR has been used as an objective measure of image quality over many years, but it correlates weakly with human quality classifications (Yao et al., 2005). EN are a measure of reconstructed brain surface complexity calculated by FreeSurfer, and has been found to correlate with movement artifacts (Rosen et al., 2018). Another option for automated QC is the exclusion of cases according to outliers based on global morphological estimates like mean cortical thickness, total surface area or estimated total intracranial volume, a technique commonly used in neuroimaging studies, e.g. (Boedhoe et al., 2018; Guadalupe et al., 2014; Shinn et al., 2017).

Additionally, several tools for a more extensive QC are currently available. Two promising tools are MRIQC (Esteban et al., 2017) and Qoala-T (Klapwijk et al., 2019). Both tools use machine learning to provide a rating for quality. While MRIQC uses the T1 or T2 images as input to provide a score on image quality, Qoala-T uses FreeSurfer’s segmentation and parcellation output, together with FreeSurfer’s output quality metrics to provide a score on segmentation quality.

The aims of this study are: 1) to determine the effect that manual editing of FreeSurfer’s output has on the resulting morphological estimates, and 2) to identify which QC approach is best in terms of reduction of unexplained variance relative to total variance in the context of large-population imaging. As a test case to assess this reduction in unexplained variance, we use a regression of morphological brain measures against age. The time investment of each QC strategy is taken into consideration, and we provide a recommendation for the optimal QC approach in large structural neuroimaging studies.

## 2 Material and Methods

### 2.1 Study design and participants

#### 2.1.1 Sample

We used data from The Maastricht Study, an observational prospective population-based cohort study (Schram et al., 2014). In brief, the study focuses on the etiology, pathophysiology, complications, and comorbidities of type 2 diabetes and is characterized by an extensive phenotyping approach. All individuals aged between 40 and 75 years living in the southern part of the Netherlands were eligible for participation. Participants were recruited through mass media campaigns, from the municipal registries and the regional Diabetes Patient Registry via mailings. Recruitment was stratified according to known type 2 diabetes status, with an oversampling of individuals with type 2 diabetes. The study has been approved by the institutional medical ethical committee (NL31329.068.10) and the Dutch Ministry of Health, Welfare, and Sports of the Netherlands (permit 131088-105234-PG). All participants gave written informed consent.

The present report uses cross-sectional data from the first 3451 participants who completed the baseline survey between November 2010 and September 2013. The selection includes 200 participants with mild cognitive impairment (MCI), and 200 non-MCI participants in order to introduce some heterogeneity in the sample. These are matched on age, sex and educational level, without oversampling for diabetes. MCI diagnosis was based on: Mini-Mental State Examination (Folstein et al., 1983) scores below 24 points; more than two cognitive tests not executed; delayed recall and word learning test (Walton, 1958), or Stroop-III (Stroop, 1935) 1.5 SD below the population-mean. Of those 400 participants, 260 had MRI brain data available. Data processing and extraction failed in one participant, specifically in the MRIQC tool processing, and was removed. Hence, the current manuscript includes 259 participants. See participant inclusion flowchart in Figure 1.

**Figure 1:**
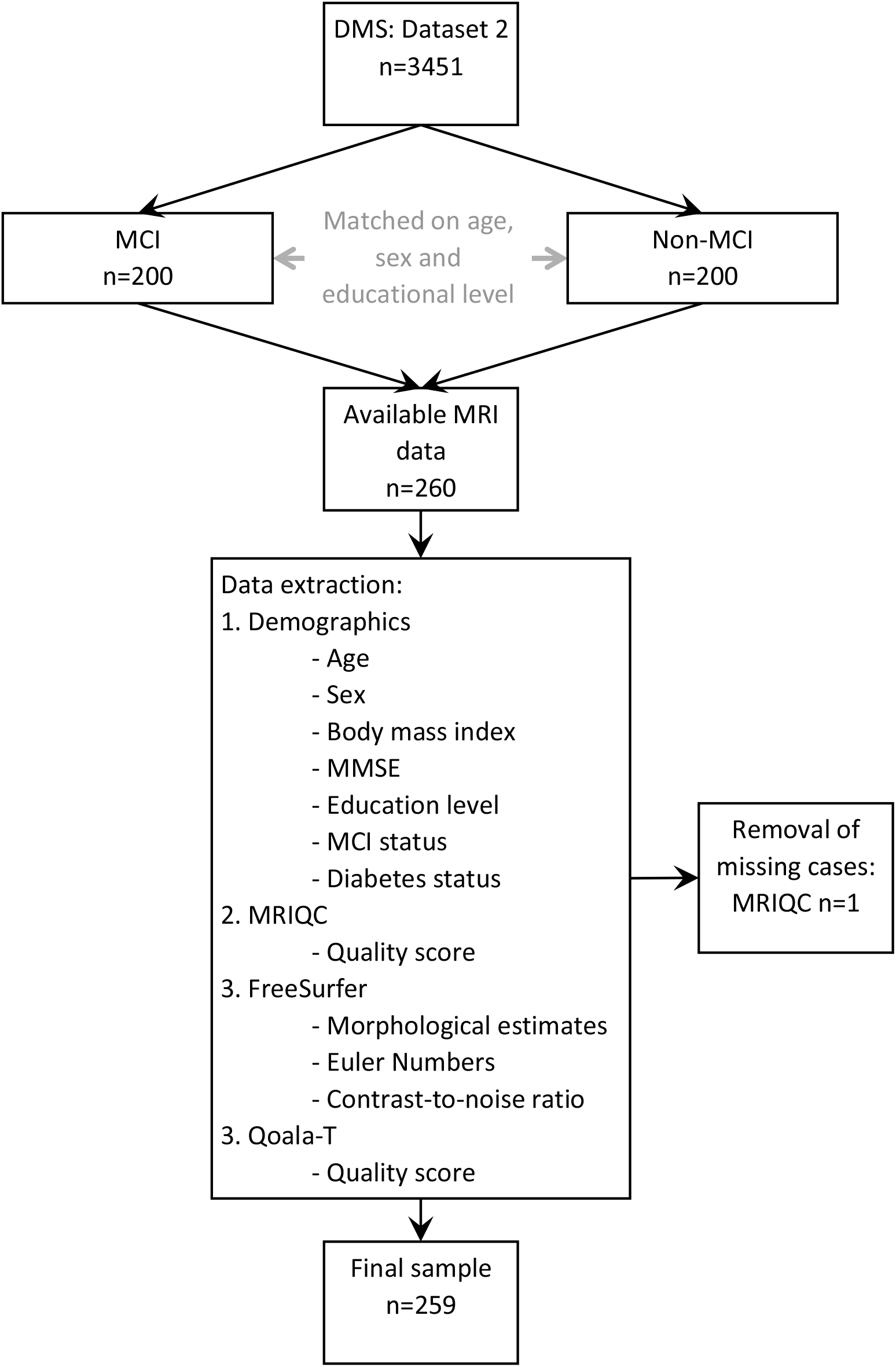
Sample selection and data extraction. Among the included participants in dataset 2, a sample of MCI participants and matched non-MCI were selected. Among these, 260 participants had available brain MRI data. There was one missing case because the MRIQC tool was unable to run in one of the participants. The case with missing data was removed and the present report includes 259 subjects. *Abbreviations:* MCI: Mild cognitive impairment; MMSE: Minimental State Examination.

#### 2.1.2 MRI acquisition

Brain images were acquired on a 3T clinical magnetic resonance scanner (MAGNETOM Prismafit, Siemens Healthineers GmbH) located at a dedicated scanning facility (Scannexus, Maastricht, The Netherlands) using a head/neck coil with 64 elements for parallel imaging. The MRI protocol included a three-dimensional (3D) T1-weighted (T1w) magnetization prepared rapid acquisition gradient echo (MPRAGE) sequence (repetition time/inversion time/echo time (TR/TI/TE) 2,300/900/2.98ms, 176 slices, 256 × 240 matrix size, 1.0 mm cubic reconstructed voxel size); and a fluid-attenuated inversion recovery (FLAIR) sequence (TR/TI/TE 5,000/1,800/394 ms, 176 slices, 512 × 512 matrix size, 0.49 × 0.49 × 1.0 mm reconstructed voxel size).

#### 2.1.3 Brain segmentation

Brain segmentation and cortical parcellation was performed on 259 participants with FreeSurfer v6.0 (Fischl, 2012) using T1w and FLAIR images as input. The optional arguments “-FLAIRpial” and “-3T” were used to optimize segmentation and parcellation quality. In addition, hippocampal subfields (Iglesias et al., 2015) were extracted. FreeSurfer output yielded cortical area (68 ROIs) and cortical thickness estimates (68 ROIs) in accordance with the Desikan-Killiany atlas (Desikan et al., 2006), as well as subcortical volumes (38 ROIs), and hippocampal subfields (24 ROIs). Hence, a total of 198 morphological estimates were obtained per individual brain. With no further manipulation, tabulated data was extracted. This original dataset will be referred from now on as “Non-QC dataset” (see Figure 2A) and used as reference for comparison with the other QC datasets.

**Figure 2:**
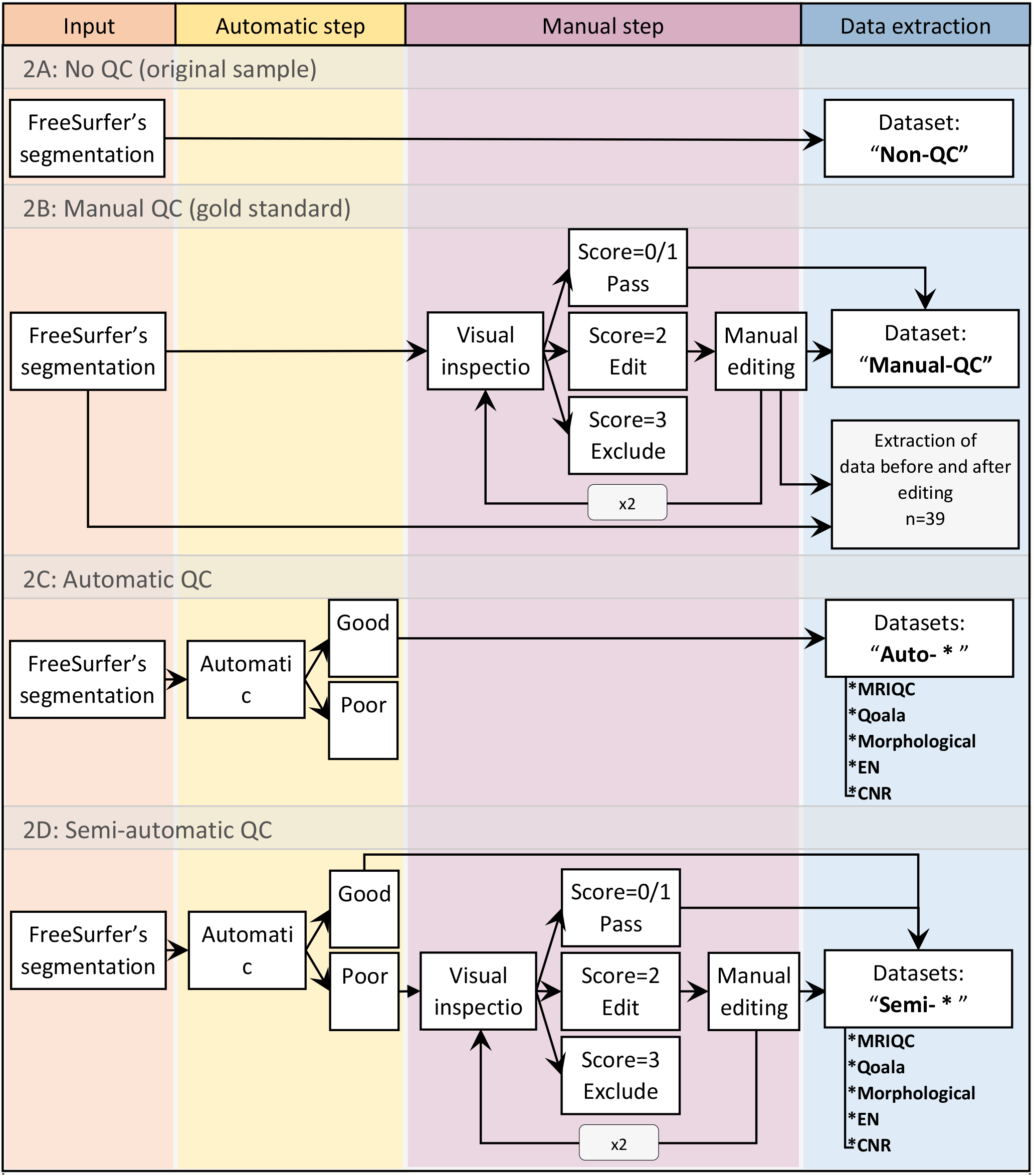
Creation of datasets through QC strategies. Each QC strategy includes specific steps: Manual QC includes only the manual step, Auto-QC includes the automatic step, Semi-QC includes both, and no QC includes none of them. <u>2A) No QC:</u>The extraction of the morphological estimates is done without any QC. <u>2B) Manual QC:</u>Segmented images undergo visual inspection. Concurred rating classifies the images to “pass”, “edit” or “exclude”. When appropriate, manual editing is performed a maximum of two times. All images classified as “pass” in the last round are included in the dataset. In parallel, data is independently extracted (box in gray) for only those images manually edited (n=39). This is done before and after undergoing manual editing. <u>2C) Automatic QC:</u>Segmentations are classified according to their quality by either an automated software (MRIQC & Qoala-T) or outliers based on several metrics (Morphological, EN or CNR). Only those classified as “good” are included in the dataset. <u>2D) Semi-automatic QC:</u>A “combination” between automatic and manual QC. Instead of being excluded, images classified as “poor” by the automated steps, undergo visual inspection and manual editing when appropriate. *Abbreviations:* QC: Quality control; Qoala: Qoala-T; EN: Euler numbers; CNR: Contrast-to-noise ratio

#### 2.1.4 Quality control strategies

Eleven QC strategies were applied to the original sample, generating 11 new QC datasets covering all morphological estimates for cortical thickness, cortical area, subcortical volumes and hippocampal volumes. Each QC strategy resulted in different brains to edit, include and exclude. Hence, all the 11 new QC datasets (and the non-QC dataset) contain 198 morphological estimates, but differ with respect to participant inclusion and which brains underwent manual editing.

These strategies can be divided into three categories: 1) full manual QC with editing, i.e. gold standard; 2) automated QC by exclusion of outliers based on: MRIQC, Qoala-T, Morphological, EN or CNR measures; 3) semi-automated QC by visual inspection of outliers based on: MRIQC, Qoala-T, Morphological, EN or CNR measures. In the next sections we describe the QC strategies in detail.

##### 2.1.4.1 Manual quality control and editing: gold standard assessment

The first category, “full manual QC” includes only one QC strategy: the gold standard. Figure 2B shows the manual QC process. This strategy was executed by two researchers who independently visually inspected and rated brain segmentations according to their quality, followed by manual editing of segmentations identified as inaccurate.

A standard operating procedure (SOP) for visual inspection and manual editing was designed specifically for The Maastricht Study. A researcher with three years of experience in hands-on QC of large MRI cohorts (J.M.) performed visual inspection of the 259 brain segmentations twice with a 6-month period gap, without and with the help of the SOP respectively. A second researcher (M.B.), without prior experience in QC, independently reviewed the same set of segmentations once, after training by rater 1, and following the same SOP. Both researchers scored the quality of the segmentations from 0 to 3, where 0 referred to segmentations with perfect quality, 1 to segmentations with sufficient quality, 2 to segmentations that needed manual editing, and 3 to segmentations that should be excluded due to unfixable inaccuracies.

Finally, both researchers met to review and discuss each discordant case and reached consensus, creating a final agreed-upon score called from now on “Accorded rating”. The segmentations with accorded ratings of 0 or 1 were accepted, and those rated with a 3 were removed from the dataset.

Subsequently, manual editing was performed on those brains with inaccurate segmentations -scored as 2 by the accorded rating-. The editing was performed by changing the brain surfaces where inaccuracies were detected. This process was done through addition or removal of voxels in the white matter mask, removal of voxels in the brain mask, and addition of control points in the brain mask.

The edited subjects subsequently underwent a new segmentation pipeline. This process-visual inspection, manual editing, and production of a new segmentation-was repeated a maximum of two times when necessary, after which, the reconstructed images were visually inspected one last time. These were then accepted as accurate (scores 0 or 1) or rejected as unfixable (score of 3).

Tabulated data from all accepted segmentations’ cortical thickness, cortical surface area, subcortical volumes and hippocampal volumes -either edited or unedited-were subsequently extracted. This tabulated data will be referred to as “Manual-QC dataset”.

In order to study changes produced by manual editing in brain estimates, morphological estimates were additionally extracted before and after manual editing across all the edited segmentations (n=39). These 39 subjects will be used to investigate the alterations due to manual editing on morphological FreeSurfer estimates.

##### 2.1.4.2 Automatic quality control: exclusion of cases

Figure 2C shows the automatic QC process. MR images were either accepted or excluded based on the assessment by the following tools: MRIQC (Esteban et al., 2017), and Qoala-T (alternative B) (Klapwijk et al., 2019); and the following metrics: FreeSurfer’s global morphological measures, EN (Dale et al., 1999), and CNR (Welvaert and Rosseel, 2013).

The tools MRIQC and Qoala-T use machine learning to provide a binary quality score (“good” or “poor”). Segmentations with a quality score of “poor” were excluded. Tabulated data were then extracted, creating the datasets “Auto-MRIQC dataset” and “Auto-Qoala dataset” respectively. Despite the fact that many studies define outliers based on standard deviation, it is a measure highly dependent on distribution, as it assumes a normal distribution, and hence not a robust method to detect outliers (Leys et al., 2013). For this reason, in this study outliers were defined as 1.5 interquartile range (IQR) below the first quartile (Q1), and 1.5 IQR above the third quartile, following the classical method proposed by Tukey (1977). Hence, the lower inner fence was defined as Q1-1.5*IQR, while the upper inner fence was Q3+1.5*IQR.

The identification of morphological outliers was specific for each type of measure, and based on the next FreeSurfer’s global estimates: left/right hemisphere mean thickness, for estimates of cortical thickness; left/right hemisphere white surface area for cortical area; estimated total intracranial volume and mask volume for subcortical volumes; and left/right hemisphere whole hippocampus volume for hippocampal subfields. Outliers were excluded below the lower and above the upper inner fences. Tabulated data were extracted for each type of morphological estimate separately (cortical thickness, cortical area, subcortical volumes and hippocampal volumes), and then put together, creating the dataset “Auto-morphological dataset”.

Outliers based on EN and CNR metrics were excluded only as values below the lower inner fence, because high values in EN and CNR indicate a positive relation with quality. Tabulated data were extracted, creating the datasets “Auto-EN dataset” and “Auto-CNR dataset” respectively.

##### 2.1.4.3 Semi-automated quality control: automatic detection with visual inspection and editing

Figure 2D shows the semi-automated QC process. Based on the same principle as for the automatic QC strategies, potentially inaccurate cases and outliers were identified with MRIQC, Qoala-T, global morphological estimates, EN, and CNR. Rather than being excluded, the potentially inaccurate cases went through visual inspection and manual editing when necessary, in an identical scheme as the one described in section “2.1.4.1 Manual quality control and editing: the gold standard assessment”.

For each approach, tabulated data were then extracted creating the QC datasets: “Semi-MRIQC dataset”, “Semi-Qoala dataset”, “Semi-Morphological dataset”, “Semi-EN dataset”, and “Semi-CNR dataset”.

### 2.2 Statistical analysis

#### 2.2.1 Sample characteristics

Wilcoxon signed-rank tests and Chi-squared (χ^2^) tests, for continuous and categorical variables respectively, were performed to assess significant differences between the MCI and non-MCI groups.

#### 2.2.2 Agreement and overlap of manual ratings

Weighted Cohen’s Kappa (К) (Cohen, 1968) and percentage of agreement 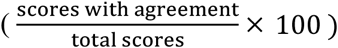 were implemented to assess inter- and intra-rater reliability among the visual inspection’s ratings.

#### 2.2.3 Manual editing effects

To assess our first aim, the percentage of change on brain morphological estimates 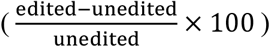 after manually editing was extracted for each of the 198 brain morphological estimates separately. Wilcoxon signed-rank test, and effect size (r) (Rosenthal et al., 1994) defined as r=Z/√N, where Z is the Z-score, and N is the sample size, were used to test significance of changes before and after manual editing for each paired morphological estimate. False discovery rate (FDR) was used for multiple comparisons correction providing q-values. In addition, we extracted the average within-subject coefficient of variation (CoV) for each of the 198 morphological estimates as:

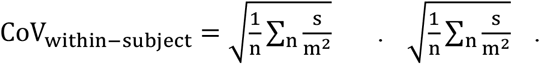

Where n is the sample size, s the within-subject variance as 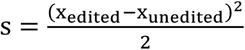, m the, within-subject mean as 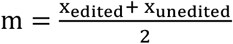, and x is a specific brain morphological estimate.

#### 2.2.4 Comparison of QC strategies

To assess our second aim, we focus on how QC strategies change the proportion of unexplained variance relative to its total variance. The background concept is based on the measurement error contained in a linear regression 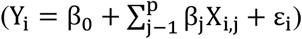 model’s stochastic component, the error term. Purely measuring the changes in unexplained variance is insufficient, as any QC strategy that would reduce the total variance of a sample will collaterally reduce the unexplained variance, but potentially also the explained one. Therefore, in this paper we will use the inverse proportion of unexplained-to-total variance. The coefficient of determination (R^2^) captures this proportion 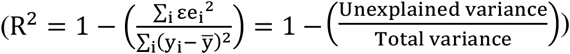. It can be robustly extracted from a linear regression model, and in practical terms it is not affected by sample size (Hayes, 2017).

In order to extract R^2^ for each morphological measure in each dataset, we created a test case regression model with one morphological measure as dependent variable, and age and sex as independent variables. We ran this model separately for each of the 198 morphological measures, in the non-QC dataset and in each of the 11 newly created datasets, obtaining 198 R^2^ for each of QC datasets (i.e. a total of 2376 R^2^ values). The reason to use a model with age and sex is to ensure a wide range of R^2^ values in each dataset. Age related brain atrophy has been widely studied (Gur et al., 1991; Kakimoto et al., 2016; Murphy et al., 1992; Tang et al., 2001; Yoshii et al., 1988), and most brain regions are to some extent affected by age. Please note that purpose of the linear regression with age is to obtain a metric related to the unexplained-to-total variance, not to assess what morphological estimate has the strongest relationship with age. Using non-QC as baseline, we then individually subtracted the R^2^ values obtained in the non-QC sample from their paired R^2^ values obtained in each of the 11 QC datasets, obtaining 198 delta R^2^ (ΔR^2^) for each of the 11 QC strategies (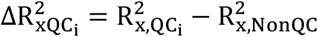, where x is the specific morphological estimate, and i corresponds to 1 of the 11 QC datasets not including the baseline, Non-QC).

An increase in 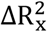 will then indicate a reduction of unexplained-to-total variance ratio for a specific brain morphological estimate, and hence a beneficial increase of relative explained variance. Differences in 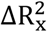 will be assessed qualitatively.

All statistical analysis were performed in R 4.0.2 (2020-06-22) (Team, 2013). Graphs were created through ggplot2 (Wickam, 2009), and brain maps through ggseg (Mowinckel & Vidal-Piñeiro, 2019). Scripts are available upon reasonable request from the corresponding authors.

## 3 Results

### 3.1 Sample characteristics

#### 3.1.1 Subjects

259 participants completed FreeSurfer’s recon-all and underwent all QC strategies. Supplementary Table 1 summarizes the characteristics of the study sample stratified for MCI and non-MCI. The MCI and non-MCI participants were matched for age, sex and educational level. A significantly higher BMI in MCI (p=0.017) was found, but there were no other differences on demographic parameters between participants.

#### 3.1.2 Brain segmentation & creation of datasets

A total of 11 QC strategies were applied to the original sample (non-QC dataset). Each strategy differed in the amount of segmentations that were visually inspected, edited or excluded, as well as the time investment of performing each QC strategy. The sample sizes of the newly created QC datasets ranged from n=205 to n=259. The time investment to perform QC ranged from 15 minutes for the auto-QC strategies to 126 hours (approximately 30 minutes per subject) when applying the manual QC strategy. Supplementary Table 2 summarizes the number of segmentations visually inspected, edited and excluded, the total sample sizes of the new datasets, and the time investment of applying each QC strategy.

#### 3.1.3 Manual quality control

The inter-rater agreement was 54.7%. The intra-rater agreement was 40.7% when only one rater used SOP, and 47.0% when both raters followed the SOP. Table 1 summarizes inter/intra rater reliability through weighted Cohen’s Kappa values (К), which takes in consideration the scores as ordered values, and confidence intervals (CI). The inter-rater reliability increases with the use of a SOP, reaching a К=0.25, similar to the intra-rater reliability, with К=0.24.

**Table 1:** Cohen’s kappa scores between raters and confidence interval. TP: time point; SOP: Standard operating procedure.

| Kappa<br>(Confidence<br>Interval) | Rater<br>1 TP1 | Rater 1<br>TP2 | Rater 2 |
| --- | --- | --- | --- |
| **Rater 1**<br>**TP1** | 1 | 0.24<br>(-0.45, 0.94) | 0.16<br>(-0.41, 0.73) |
| **Rater 1**<br>**TP2** | – | 1 | 0.25<br>(-0.09, 0.60) |
| **Rater 2** | – | – | 1 |

The accorded rating, agreed-upon by both raters, classified 17.8% of the segmentations as inaccurate, either by requiring manual editing or exclusion. Figure 3 shows the distribution of the scores given by each rater, as well as the final accorded rating.

**Figure 3:**
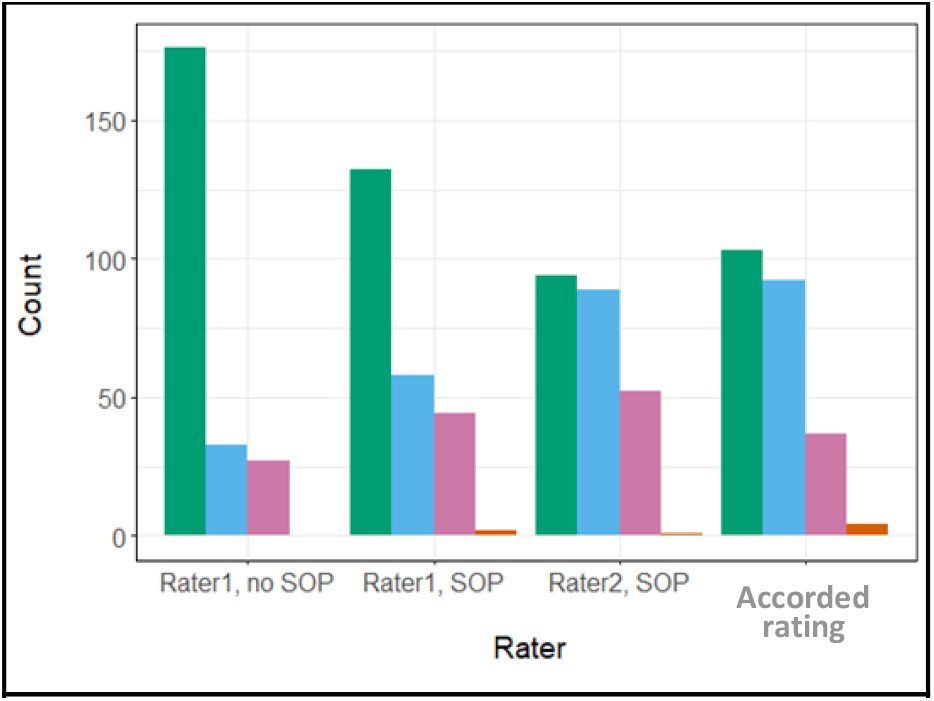
Distribution of given scores in each visual inspection and final accorded rating.

### 3.2 Manual editing of brain surfaces: changes in brain estimates

Segmentations of 39 out of 259 brains had an accorded rating of “2” and thus were manually edited. Manual editing resulted in changes in all morphological measures out of the 39 edited brains. The largest difference after editing, with a mean increase of 25% of its volume, was found in bilateral fimbria, followed by differences that ranged from +9 to +12% in bilateral vessel and cerebellar white matter. The largest reduction was found in the fourth ventricle, with a mean volume reduction of 7%. Figure 4A shows the average difference in percentage across subjects after editing.

**Figure 4:**
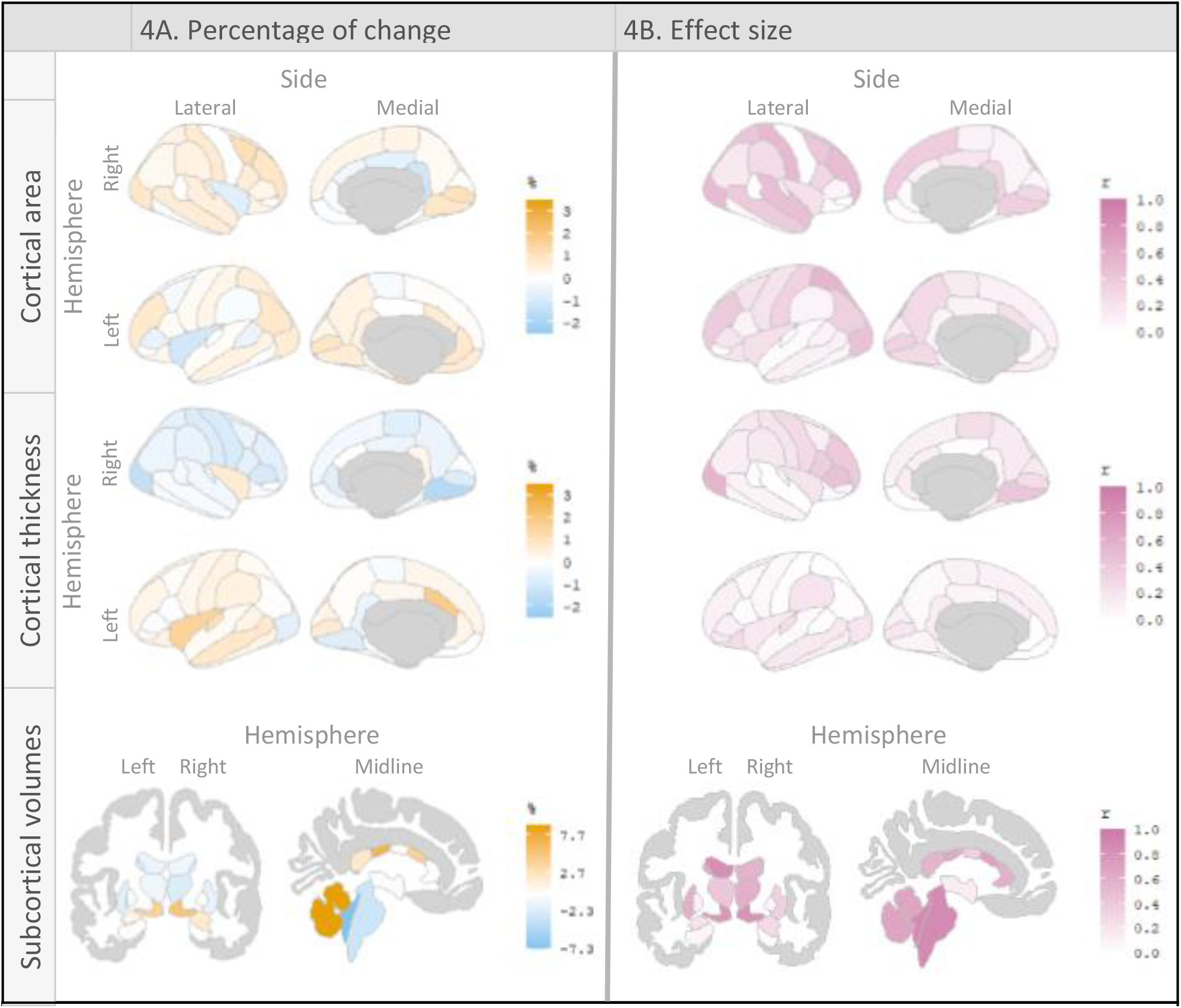
Brain maps. 4A: Figure shows the average change after editing of each ROI. Yellow color indicates an increase in volume, area or thickness, while blue color shows a decrease. Color intensity indicated magnitude. Notice the percentage of change in subcortical volumes is larger than those found in cortical areas and thickness, and so it is reflexed it its legend. 4B: Effect size. Brain map shows the effect size (r) of the changes, larger effect sizes are closer to 1 and represented in pink, while lower effect sizes are closer to 0 and represented in white. *Note: The brain maps do not represent all the analyzed morphological measures, for a list of percentage of change, q-values and effect sizes of all brain regions see Supplementary Table 3 (A, B and C)*.

Wilcoxon signed-rank test showed significant differences (q-value < 0.05) in 53 out of the 197 analyzed morphological measures. See Supplementary Figure 1 for q-values’ brain maps. The effect sizes (r) ranged from 0 to 0.87, with the largest effect sizes (r>0.8) found in bilateral fimbria, brainstem, 4th ventricle, left cerebellum white matter, left lateral ventricle, and bilateral ventral DC. Figure 4B shows the effect size distribution for several cortical and subcortical morphological measures. A brain map legend is provided in Supplementary Figure 2.

The median CoV for all brain regions was 2.4%, with a standard deviation (SD) of 3.0%. The largest CoV was found within subcortical structures (CoV=4.5%, SD=4.6%), followed by hippocampal subfields (CoV=2.9%, SD=4.2%), cortical thickness (CoV=2.2%, SD=1.0%), and the smaller in cortical areas (CoV=1.7%, SD=1.6%).

### 3.3 Quality control Approaches: consequences in a regression analysis

R2 was extracted for 198 morphological measure over 12 datasets (1 non-QC and 11 QC datasets). The distribution of the R2 values obtained by each dataset can be found in Supplementary Figure 3.

Figure 5 shows the mean ΔR^2 obtained by each QC strategy, when compared to the non QC dataset. The QC strategies that resulted in a higher increase in explained to total variance ratio as measured by larger positive ΔR^2 are: Manual QC (mean ΔR^2=.0087) for cortical thickness; Auto Qoala (mean ΔR^2=.0115) and Auto EN (mean ΔR^2=.0077) for cortical area; auto MRIQC (mean ΔR^2=.0123), Auto EN (mean ΔR^2=.0097), and Auto Qoala (mean ΔR^2=.0060) for subcortical volumes; and Auto EN (mean ΔR^2=.0131), Auto MRIQC (mean ΔR^2=.0072) and Auto Qoala (mean ΔR^2=.0055) for hippocampal subfields. The exclusion of cases based on morphological estimates (Auto Morphological) produces a relatively large reduction in R2 for any type of measure (mean ΔR^2= .0337 for area, mean ΔR^2= .0222 for hippocampal subfields, mean ΔR^2= .0172 for subcortical volumes and mean ΔR^2= .0107 for cortical thickness), as well as Auto Qoala, which reduces R2 in cortical thickness in .0207 points relative to the non QC strategy.

**Figure 5:**
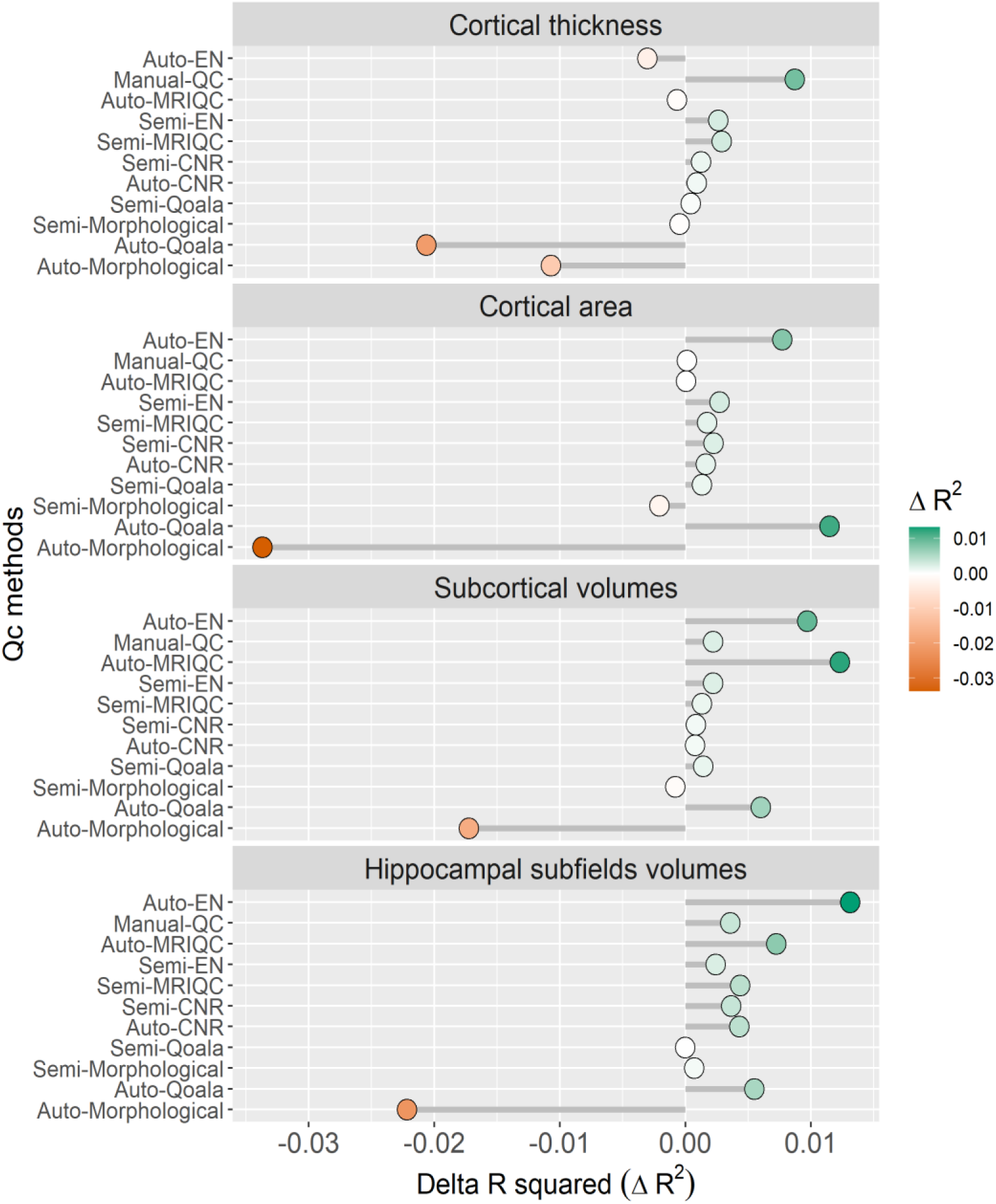
Circles position indicate the mean Δ*R*^2^ across type of morphological estimates obtained by each QC strategy. Green indicates an increase in R^2^ while orange indicates a decrease, intensity of color shows magnitude of change.

Across all brain regions, the exclusion based on Euler numbers (Auto-EN) yields the largest decrease of unexplained variance relative to total variance, as shown by the largest increase in R^2^ (Δ*R*^2^ =.0050). Auto-Morphological yields the largest decrease in R^2^ (Δ*R*^2^ =.-.0213). Figure 6 show the mean Δ*R*^2^ for each QC strategy in each type of morphological measure.

**Figure 6:**
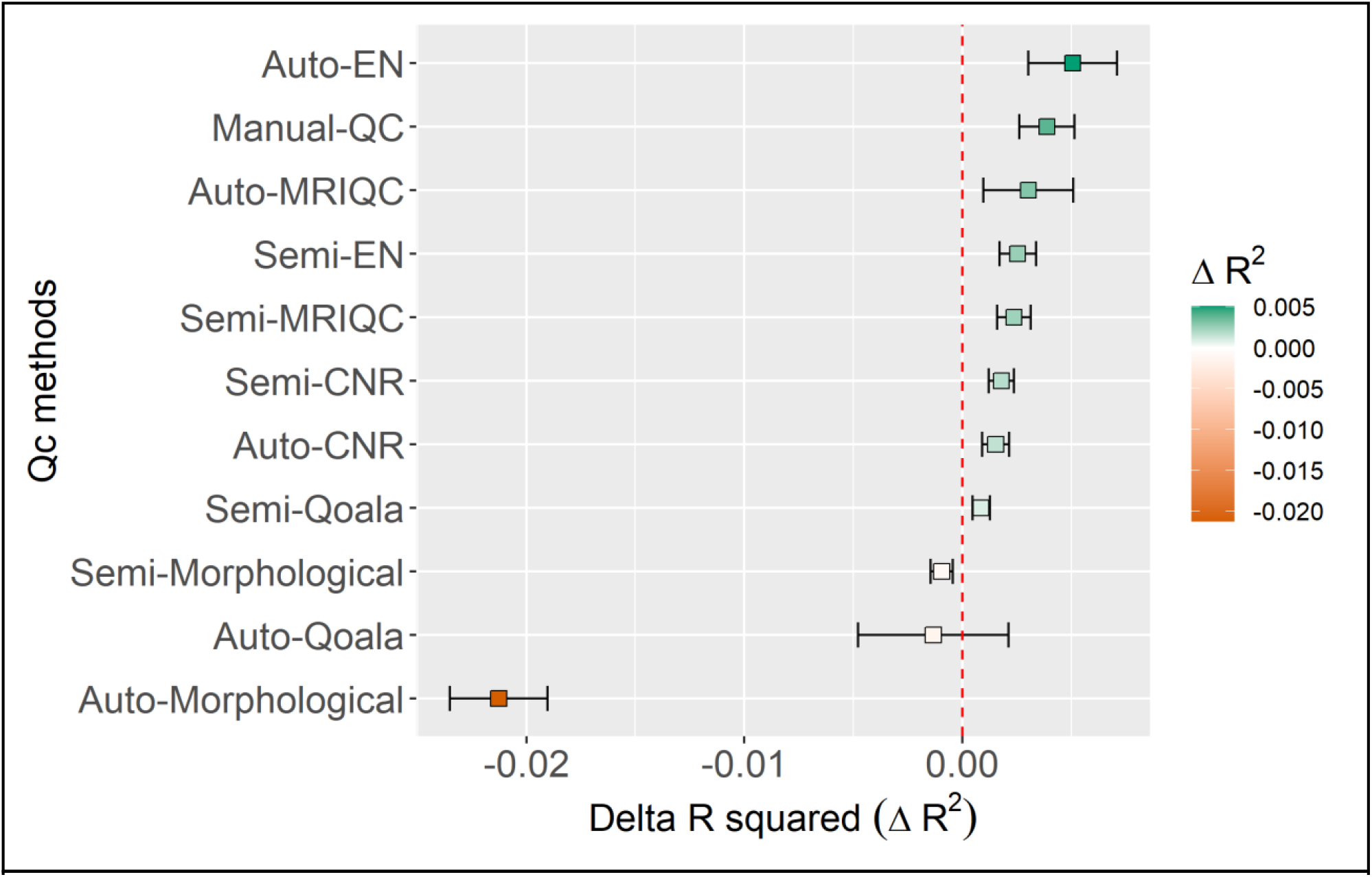
Squares indicate the mean Δ*R*^2^ across morphological estimates obtained by each QC strategy. Whiskers show the 95% confidence interval for the mean. Green indicates an increase in R^2^ while orange indicates a decrease, intensity of color shows magnitude of change. Red dotted line shows the mean Δ*R*^2^ of the non-QC strategy, i.e. zero. *Abbreviations: QC: Quality control; EN: Euler numbers; CNR: Contrast-to-noise ratio; Qoala: Qoala-T*

## 4 Discussion

In this study we investigated the influence of manual editing of brain surfaces on resulting morphological FreeSurfer estimates. We also compared the performance of different QC strategies by assessing how these alter the relative proportion of unexplained variance in a neuroimaging sample.

### 4.1 Manual editing brain surfaces

Manual editing of brain segmentations produces significant changes in several brain morphological estimates across thickness, surface area and volumetric measures, with large effect sizes especially found in subcortical structures. Similar studies showed mixed results: Waters et al. (2019) showed significant differences in cortical but not subcortical volume following editing. Beelen et al. (2020) found a significant increase in cortical area, and a decrease in cortical thickness after manual editing. Conversely, McCarthy et al. (2015) found no significant differences in cortical area, thickness, and subcortical volumes. The discrepancy in results could be explained by the use of different editing techniques: While Waters et al. (2019) only edited voxels in the brain mask, Beelen et al. (2020) also edited the white matter mask, and used control points; McCarthy et al. (2015) exclusively used control points and white matter modification; and we edited the brain mask, white matter mask and used control points.

The within-subject CoV that we obtained after manual editing was similar to what was found in repeatability studies for subcortical structures (Maclaren et al., 2014; Velasco-Annis et al., 2018). Further, similar CoV were found in FreeSurfer’s reproducibility studies between operating systems in measures of volume and thickness (Gronenschild et al., 2012). Taken together, our results indicate that despite the significant changes in morphological estimates produced by manual editing, those changes are similar to those expected from the process of re-running the segmentation process itself, and do not necessarily imply an improvement in segmentation quality.

### 4.2 Comparison between QC methods

We created a regression model and extracted the ΔR^2^ for each morphological estimate in each QC strategy when compared with non-QC. To our knowledge, this is the first study using the proportion of unexplained-to-total variance as a measure of quality.

Two previous studies have assessed the importance of manual QC by testing whether it increases the sensitivity to detect differences between clinical groups. Both studies used paired t-tests to compare the effect sizes obtained with and without QC, with no significant results (Beelen et al., 2020; McCarthy et al., 2015). With a different methodology, Waters et al. (2019) assessed if the correlation coefficients for brain-behavior relationships differed between edited and unedited segmentations, finding non-significant differences. Relating the quality of the data to the capacity to find significant results can be misleading, as non-randomized noise, for example the one caused by biases in the data (Blumenthal et al., 2002; Madan, 2018; Pardoe et al., 2016; Savalia et al., 2017; Wenger et al., 2014) can lead to more significant differences between groups. Similarly, some studies have used the variance of brain volumes before and after applying a specific QC strategy (Backhausen et al., 2016; McCarthy et al., 2015) to test whether a QC strategy is adequate. And while the reduction of the total variance of a sample implies a potential reduction on unexplained variance, it can also entail a potential reduction of explained variance driven by actual morpho-physiological information. Finally, other studies assess the viability of a QC method by comparing it to manual ratings (Klapwijk et al., 2019; Rosen et al., 2018; Yao et al., 2005), which assumes that manual QC improves the quality of a sample. Using the proportion of unexplained-to-total variance, allows us to assess the measurement error caused by both noise and biases.

#### 4.2.1 The optimal QC strategy depends on the type of morphological estimate of interest

Our results show that cortical thickness benefits the most from manual QC, despite the small changes by manual editing in cortical thickness; cortical area benefits most from Auto-Qoala and Auto-EN; and subcortical and hippocampal subfields volumes from Auto-EN and Auto-MRIQC.

The different effects of the QC strategies on each type of estimate can be explained by the particular mechanisms of each QC strategy. Manual-QC visually inspects the upper and lower boundaries of the cortical gray matter (pial and white matter surfaces respectively), which are tightly related to the cortical gray matter thickness. Manual editing changes the distance between these boundaries, directly changing the cortical thickness estimates. However, manual editing does not change the boundaries between specific ROIs, and hence morphological estimates of surface area are not substantially affected. Distinct from cortical parcellation, subcortical segmentation is defined based on image intensities and probabilistic information of ROIs positions (Fischl et al., 2002), and thus, correcting errors in surface boundaries and adding control points to correct normalization errors changes the subcortical estimations only indirectly. The QC strategies based on image quality metrics -such as CNR and specially MRIQC-may provide a good indication of segmentation quality based on the fact that good image quality -for example high contrast, with no movement or artifacts-which facilitates good segmentation performance.

#### 4.2.2 QC strategies based on global morphological estimates are not a suitable QC solution

Notably, for all types of measures, the ΔR^2^ decreased by the commonly used strategy of excluding or visually inspecting and editing subjects based on global morphological estimates (Auto- and Semi-Morphological QC strategies). Excluding outliers based on morphological estimates naturally reduces the total variance of a sample, but our results indicate that a large part of this reduced variance is not unexplained variance but potentially relevant morpho-physiological information. In addition, a previous study found that approximately 40% of the segmentation errors are not identified by the use of morphological outliers (Waters et al., 2019). Taken together, this strongly indicates that the exclusion of subjects based on global morphological estimates is not a suitable QC strategy.

#### 4.2.3 Auto-EN produces, on average, the greatest reduction in noise

Some studies might be interested in all types of morphological measures at the same time. For those cases, Auto-EN is a pragmatic strategy that (on average) produces the best results, consistently good for three out of four types of brain measures (i.e. subcortical volumes, hippocampal subfields and cortical area). EN are a measure of reconstructed surface complexity, and has been found to highly correlate with visual inspection scores and image artifacts in several samples (Rosen et al., 2018). It is perhaps counterintuitive that this QC strategy performs poorly for measures of cortical thickness. However, EN relates to the frequency, and not the size, of bridges and holes on the brain segmentation. Cortical thickness might be influenced by large errors (possibly more related to skull striping) than from frequent small ones, which may be more related to underlying image quality. Hence, auto-EN is an effective and time-efficient QC strategy despite the small reduction in ΔR^2^ for cortical measures.

Alternatively, manual-QC and semi-EN provide a small but consistently good increase in ΔR^2^, all the same with a higher time investment.

### 4.3 Limitations

The design of this study does not allow drawing firm conclusions beyond models using age as an independent variable. However, a large proportion of published studies include age as a covariate or independent variable, and thus, information about the optimal QC procedure for such studies will be relevant for the neuroimaging community. In addition, the segmentation quality of our sample was high, as measured by a high percentage (82.2%) of brain segmentations rated as accurate. Different (more severely affected) clinical samples, studies with different population types, or even different acquisition hardware or parameters, could lead to a different pattern of segmentation inaccuracies, and further research needs to be performed to investigate these scenarios.

## 5 Conclusion

Manual editing of brain surfaces significantly alters, but not necessarily improves, FreeSurfer’s brain morphological estimates. The selection of a QC strategy should be determined by the type of morphological measures of interest in a study, while taking in consideration the available resources.

We recommend the exclusion of outliers based on Euler number for studies using subcortical measures, hippocampal subfields or cortical areas, as it produces on average the largest increase in explained-to-total variance proportion, in addition to being a highly time-efficient QC strategy. Manual-QC is the strategy that may provide the best quality for studies including cortical thickness measures. However, this strategy may not be feasible for large samples, and the exclusion of outliers based on Euler Numbers or MRIQC, or the visual inspection of outliers based on Euler numbers are reliable alternatives. Finally, we discourage the exclusion of outliers based on global morphological estimates as a method to ensure segmentation quality as it reduces a larger proportion of explained variance.

## Acknowledgements and founding

This study was supported by the European Regional Development Fund via OP-Zuid, the Province of Limburg, the Dutch Ministry of Economic Affairs (grant 31O.041), Stichting De Weijerhorst (Maastricht, The Netherlands), the Pearl String Initiative Diabetes (Amsterdam, The Netherlands), the Cardiovascular Center (CVC, Maastricht, the Netherlands), CARIM School for Cardiovascular Diseases (Maastricht, The Netherlands), CAPHRI Care and Public Health Research Institute (Maastricht, The Netherlands), NUTRIM School for Nutrition and Translational Research in Metabolism (Maastricht, the Netherlands), Stichting Annadal (Maastricht, The Netherlands), Health Foundation Limburg (Maastricht, The Netherlands), and by unrestricted grants from Janssen-Cilag B.V. (Tilburg, The Netherlands), Novo Nordisk Farma B.V. (Alphen aan den Rijn, the Netherlands), and Sanofi-Aventis Netherlands B.V. (Gouda, the Netherlands).

