## Supplement for "Quality control strategies for brain MRI segmentation and parcellation: practical approaches and recommendations - insights from The Maastricht Study"

### Supplementary material

#### Supplementary Table 1

Supplementary Table 1: Characteristics of the study sample (n=259) stratified by MCI and non-MCI. Wilcoxon Signed Rank Test and Chi-Square test were used to assess differences between groups. Abbreviations: MCI: Mild cognitive impairment; SD: standard deviation; BMI: Body mass index; MMSE: Mini Mental Score Examination; T2DM: Type 2 diabetes mellitus.

| | Non-MCI | MCI | Wilcox/$\chi^2$ p-value |
|---|---|---|---|
| N (count) | 127 | 132 | |
| Age (mean (SD)) | 59.45 (8.80) | 59.20 (8.61) | 0.768 |
| BMI (mean (SD)) | 25.50 (3.69) | 26.47 (4.14) | 0.013 |
| MMSE (mean (SD)) | 28.86 (1.31) | 28.46 (1.66) | 0.074 |
| Sex (% females) | 45.5 | 48.5 | 0.428 |
| Level of education (n low/medium/high) | 46/34/47 | 49/34/49 | 0.980 |
| T2DM (% yes) | 5.5 | 10.6 | 0.113 |

**Supplementary Table 2**

Supplementary Table 2: Number of segmentations visually inspected, manually edited or excluded for each QC strategy, sample sizes and time investment. The Auto and Semi-morphological QC strategies (see morphological estimate's types in italic) differ in sample size, as the including, excluding, editing criteria is based in different global estimates (see section "2.1.4.2 Automatic QC: exclusion of cases" for details). The time investment includes only the time a person has to spend on applying a specific QC method, this includes the time for visual inspection (with two independent raters), for discussion to reach the accorded rating, and the time investment of manual editing. The time investment does not include the preparation time (install the software, or prepare a pipeline, for example) nor the software running time. Abbreviations: QC: Quality control; EN: Euler numbers; CNR: Contrast-to-noise ratio.

| Category | QC strategy | Visual inspection (n) | Manual editing (n) | Exclusion (n) | Sample size (n) | Cumulative time investment (hours) |
|---|---|---|---|---|---|---|
| A. No QC | Non-QC | 0 | 0 | 0 | 259 | 0.0 |
| B. Manual QC | Manual-QC | 259 | 39 | 7 | 252 | 126.0 |
| C. Automatic QC | Auto-MRIQC | 0 | 0 | 29 | 230 | 0.1 |
| | Auto-Qoala | 0 | 0 | 54 | 205 | 0.1 |
| | Auto-morphological: | 0 | 0 | | | 0.1 |
| | Cortical thickness | 0 | 0 | 15 | 244 | |
| | Cortical area | 0 | 0 | 24 | 235 | |
| | Subcortical volumes | 0 | 0 | 23 | 236 | |
| | Hippocampal subfields | 0 | 0 | 38 | 221 | |
| | Auto-EN | 0 | 0 | 26 | 233 | 0.1 |
| | Auto-CNR | 0 | 0 | 2 | 257 | 0.1 |
| D. Semi-automatic QC | Semi-MRIQC | 29 | 8 | 2 | 257 | 16.2 |
| | Semi-Qoala | 54 | 10 | 1 | 258 | 27.5 |
| | Semi-morphological: | 67 | 7 | | | 31.3 |
| | Cortical thickness | 49 | 2 | 0 | 259 | |
| | Cortical area | 38 | 3 | 1 | 258 | |
| | Subcortical volumes | 31 | 3 | 1 | 258 | |
| | Hippocampal subfields | 58 | 3 | 2 | 257 | |
| | Semi-EN | 26 | 10 | 3 | 256 | 16.0 |
| | Semi-CNR | 2 | 0 | 1 | 258 | 1.0 |

#### Supplementary Figure 1

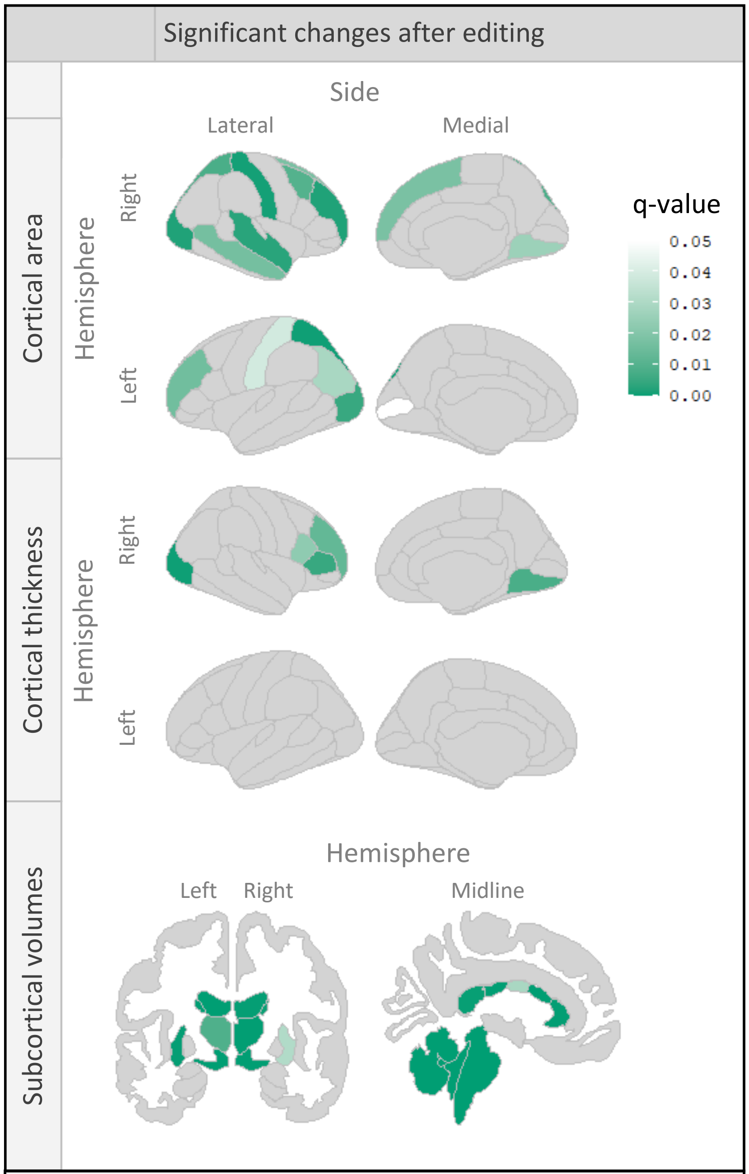


**Supplementary Figure 1**: Brain maps show in color the morphological estimates that significantly changed after manual editing.

#### Supplementary Figure 2

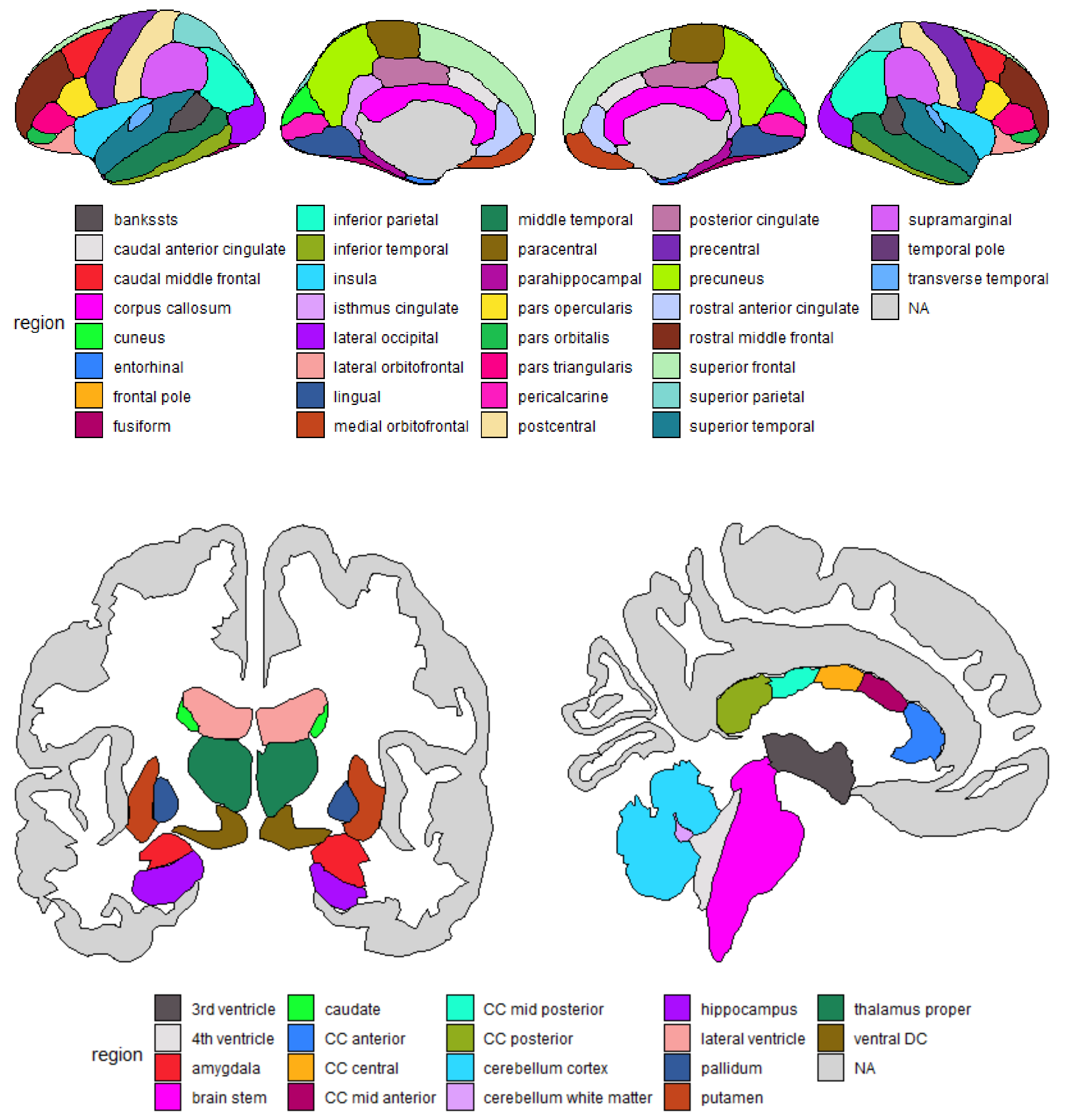


**Supplementary Figure 2:** Color coded brain maps legend with all the represented regions.

**Supplementary Table 3.A: Cortical morphological estimates**

| | Area | | | Thickness | | |
|---|---|---|---|---|---|---|
| ROI | Percentage of change (%) | q-value | Effect size (r) | Percentage of change (%) | q-value | Effect size (r) |
| LH_bankssts | 0,394 | 0,879 | 0,016 | -0,488 | 0,530 | 0,102 |
| LH_caudalanteriorcingulate | 1,854 | 0,399 | 0,136 | 0,655 | 0,157 | 0,230 |
| LH_caudalmiddlefrontal | 0,097 | 0,796 | 0,042 | -0,207 | 0,176 | 0,218 |
| LH_cuneus | 0,346 | 0,517 | 0,104 | 0,333 | 0,361 | 0,148 |
| LH_entoRH_inal | 0,442 | 0,640 | 0,076 | 1,289 | 0,498 | 0,110 |
| LH_frontalpole | -0,152 | 0,298 | 0,168 | -0,253 | 0,856 | 0,031 |
| LH_fusiform | -0,055 | 0,704 | 0,063 | 0,205 | 0,586 | 0,088 |
| LH_inferiorparietal | 0,062 | 0,911 | 0,019 | 0,795 | 0,029 | 0,350 |
| LH_inferiortemporal | 0,394 | 0,430 | 0,127 | -0,072 | 0,615 | 0,082 |
| LH_insula | 1,560 | 0,312 | 0,163 | -1,075 | 0,157 | 0,228 |
| LH_isthmuscingulate | -0,437 | 0,406 | 0,134 | 0,491 | 0,586 | 0,088 |
| LH_lateraloccipital | -0,649 | 0,233 | 0,192 | 0,624 | 0,005 | 0,451 |
| LH_lateralorbitofrontal | 0,436 | 0,120 | 0,250 | 0,285 | 0,209 | 0,202 |
| LH_lingual | -0,765 | 0,596 | 0,086 | 0,630 | 0,083 | 0,273 |
| LH_medialorbitofrontal | -0,004 | 0,967 | 0,008 | 0,676 | 0,387 | 0,140 |
| LH_middletemporal | 0,808 | 0,295 | 0,169 | 0,434 | 0,507 | 0,107 |
| LH_paracentral | -0,200 | 0,562 | 0,095 | -0,162 | 0,325 | 0,159 |
| LH_parahippocampal | -0,009 | 0,660 | 0,072 | 0,261 | 0,701 | 0,060 |
| LH_parsopercularis | -0,038 | 0,879 | 0,022 | 0,178 | 0,533 | 0,099 |
| LH_parsorbitalis | 0,199 | 0,332 | 0,156 | 0,024 | 0,994 | 0,011 |
| LH_parstriangularis | -0,121 | 0,477 | 0,115 | -0,526 | 0,238 | 0,190 |
| LH_pericalcarine | 0,685 | 0,255 | 0,183 | 0,874 | 0,050 | 0,326 |
| LH_postcentral | 0,411 | 0,972 | 0,007 | 0,356 | 0,039 | 0,332 |
| LH_posteriorcingulate | 0,283 | 0,643 | 0,077 | 0,000 | 0,556 | 0,099 |
| LH_precentral | 0,264 | 0,701 | 0,063 | 0,173 | 0,214 | 0,200 |
| LH_precuneus | 0,089 | 0,738 | 0,055 | 0,413 | 0,087 | 0,272 |
| LH_rostralanteriorcingulate | 0,356 | 0,972 | 0,007 | 0,794 | 0,759 | 0,050 |
| LH_rostralmiddlefrontal | 0,321 | 0,498 | 0,110 | 0,727 | 0,016 | 0,388 |
| LH_superiorfrontal | 0,310 | 0,494 | 0,111 | 0,098 | 0,490 | 0,112 |
| LH_superiorparietal | 0,153 | 0,989 | 0,003 | 0,582 | 0,001 | 0,544 |
| LH_superiortemporal | 0,182 | 0,754 | 0,051 | 0,102 | 0,823 | 0,037 |
| LH_supramarginal | 0,458 | 0,230 | 0,193 | -0,155 | 0,586 | 0,088 |
| LH_temporalpole | 2,111 | 0,066 | 0,295 | -0,938 | 0,519 | 0,103 |
| LH_transversetemporal | 1,116 | 0,145 | 0,235 | -0,015 | 0,670 | 0,069 |
| RH_bankssts | -0,213 | 0,851 | 0,031 | 0,041 | 0,917 | 0,018 |
| RH_caudalanteriorcingulate | -0,002 | 0,464 | 0,118 | -0,361 | 0,507 | 0,107 |
| RH_caudalmiddlefrontal | -0,460 | 0,459 | 0,120 | 1,133 | 0,010 | 0,411 |

| | | | | | | |
|---|---|---|---|---|---|---|
| RH_cuneus | -0,042 | 0,353 | 0,153 | 0,074 | 0,691 | 0,065 |
| RH_entoRH_inal | 0,523 | 0,209 | 0,203 | -1,068 | 0,277 | 0,175 |
| RH_frontalpole | -1,598 | 0,002 | 0,487 | 0,539 | 0,950 | 0,011 |
| RH_fusiform | -0,343 | 0,928 | 0,016 | -0,044 | 0,839 | 0,035 |
| RH_inferiorparietal | -0,470 | 0,430 | 0,127 | 0,259 | 0,283 | 0,169 |
| RH_inferiortemporal | -0,181 | 0,691 | 0,065 | -0,149 | 0,759 | 0,050 |
| RH_insula | 0,680 | 0,222 | 0,197 | -0,720 | 0,161 | 0,226 |
| RH_isthmuscingulate | 0,286 | 0,630 | 0,078 | -1,111 | 0,051 | 0,314 |
| RH_lateraloccipital | -1,449 | 0,001 | 0,549 | 0,844 | 0,002 | 0,508 |
| RH_lateralorbitofrontal | -0,172 | 0,764 | 0,049 | 0,349 | 0,895 | 0,022 |
| RH_lingual | -1,699 | 0,008 | 0,423 | 0,835 | 0,026 | 0,356 |
| RH_medialorbitofrontal | -0,225 | 0,954 | 0,008 | -0,068 | 0,917 | 0,018 |
| RH_middletemporal | -0,160 | 0,607 | 0,087 | 0,652 | 0,017 | 0,384 |
| RH_paracentral | -0,725 | 0,135 | 0,240 | 0,391 | 0,567 | 0,093 |
| RH_parahippocampal | -0,308 | 0,368 | 0,145 | 0,067 | 0,994 | 0,009 |
| RH_parsopercularis | -0,640 | 0,023 | 0,364 | 0,618 | 0,060 | 0,306 |
| RH_parsorbitalis | -0,091 | 0,701 | 0,063 | 0,161 | 1,000 | 0,000 |
| RH_parstriangularis | -0,940 | 0,005 | 0,455 | 0,360 | 0,630 | 0,078 |
| RH_pericalcarine | -0,799 | 0,121 | 0,249 | 1,049 | 0,060 | 0,302 |
| RH_postcentral | -0,851 | 0,267 | 0,194 | 0,690 | 0,001 | 0,515 |
| RH_posteriorcingulate | -0,453 | 0,895 | 0,022 | -0,603 | 0,258 | 0,184 |
| RH_precentral | -0,898 | 0,090 | 0,274 | -0,021 | 0,950 | 0,011 |
| RH_precuneus | -0,479 | 0,325 | 0,159 | 0,163 | 0,582 | 0,088 |
| RH_rostralanteriorcingulate | -0,724 | 0,422 | 0,130 | -0,135 | 0,829 | 0,036 |
| RH_rostralmiddlefrontal | -0,681 | 0,013 | 0,399 | 0,868 | 0,002 | 0,489 |
| RH_superiorfrontal | -0,317 | 0,516 | 0,105 | 0,341 | 0,018 | 0,379 |
| RH_superiorparietal | -0,625 | 0,315 | 0,162 | 0,383 | 0,008 | 0,424 |
| RH_superiortemporal | -0,346 | 0,913 | 0,009 | 0,515 | 0,003 | 0,474 |
| RH_supramarginal | -0,530 | 0,308 | 0,164 | 0,437 | 0,103 | 0,268 |
| RH_temporalpole | -0,436 | 0,863 | 0,029 | 0,729 | 0,336 | 0,155 |
| RH_transversetemporal | -0,464 | 0,738 | 0,055 | 0,506 | 0,593 | 0,077 |

**Supplementary Table 3.B: Subcortical morphological estimates**

| ROI | Percentage of change (%) | q-value | Effect size (r) |
|---|---|---|---|
| BrainStem | -3,106 | 0,000 | 0,863 |
| CCAnterior | 0,027 | 0,000 | 0,588 |
| CCCentral | 0,337 | 0,029 | 0,351 |
| CCMidAnterior | 4,390 | 0,000 | 0,697 |
| CCMidPosterior | 7,007 | 0,000 | 0,641 |
| CCPosterior | 1,993 | 0,000 | 0,556 |
| CSF | 1,870 | 0,096 | 0,268 |
| LH_Accumbens | 3,422 | 0,066 | 0,295 |
| LH_Amygdala | 0,907 | 0,199 | 0,208 |
| LH_Caudate | -2,168 | 0,000 | 0,632 |
| LH_CerebellumCortex | -2,189 | 0,000 | 0,749 |
| LH_CerebellumWhiteMatter | 11,036 | 0,000 | 0,838 |
| LH_Hippocampus | 0,170 | 0,694 | 0,065 |
| LH_InfLatVent | 0,484 | 0,929 | 0,016 |
| LH_LateralVentricle | -1,164 | 0,000 | 0,827 |
| LH_Pallidum | -0,439 | 0,503 | 0,109 |
| LH_Putamen | -1,766 | 0,000 | 0,589 |
| LH_ThalamusProper | -0,985 | 0,009 | 0,416 |
| LH_VentralDC | 4,168 | 0,000 | 0,778 |
| LH_choroidplexus | 4,809 | 0,005 | 0,451 |
| LH_vessel | 12,239 | 0,043 | 0,324 |
| OpticChiasm | 0,976 | 0,829 | 0,036 |
| RH_Accumbens | 3,375 | 0,008 | 0,423 |
| RH_Amygdala | 1,938 | 0,085 | 0,277 |
| RH_Caudate | -1,848 | 0,000 | 0,539 |
| RH_CerebellumCortex | -1,924 | 0,000 | 0,670 |
| RH_CerebellumWhiteMatter | 8,980 | 0,000 | 0,675 |
| RH_Hippocampus | -0,038 | 0,885 | 0,025 |
| RH_InfLatVent | 3,953 | 0,003 | 0,474 |
| RH_LateralVentricle | -0,839 | 0,000 | 0,568 |
| RH_Pallidum | 0,226 | 0,625 | 0,079 |
| RH_Putamen | -1,482 | 0,031 | 0,344 |
| RH_ThalamusProper | -2,145 | 0,000 | 0,592 |
| RH_VentralDC | 5,416 | 0,000 | 0,816 |
| RH_choroidplexus | 3,298 | 0,075 | 0,286 |
| RH_vessel | 9,065 | 0,265 | 0,181 |
| x3rdVentricle | -0,303 | 0,410 | 0,133 |
| x4thVentricle | -7,212 | 0,000 | 0,863 |

**Supplementary Table 3.C: Hippocampal subfields morphological estimates**

| ROI | Percentage of change (%) | q-value | Effect size (r) |
|---|---|---|---|
| LH_CA1 | -0,305 | 0,994 | 0,002 |
| LH_CA3 | 1,088 | 0,142 | 0,237 |
| LH_CA4 | 0,261 | 0,372 | 0,145 |
| LH_GCMLDG | 0,307 | 0,343 | 0,154 |
| LH_HATA | 1,199 | 0,194 | 0,210 |
| LH_Hippocampaltail | -2,195 | 0,001 | 0,534 |
| LH_fimbria | 24,506 | 0,000 | 0,869 |
| LH_hippocampalfissure | -0,477 | 0,194 | 0,210 |
| LH_molecularlayerHP | 1,494 | 0,010 | 0,409 |
| LH_parasubiculum | -0,309 | 0,204 | 0,206 |
| LH_presubiculum | -1,074 | 0,051 | 0,313 |
| LH_subiculum | -0,242 | 0,435 | 0,127 |
| RH_CA1 | -0,595 | 0,070 | 0,290 |
| RH_CA3 | 1,974 | 0,001 | 0,505 |
| RH_CA4 | 0,672 | 0,209 | 0,203 |
| RH_GCMLDG | 0,628 | 0,225 | 0,197 |
| RH_HATA | 1,774 | 0,704 | 0,063 |
| RH_Hippocampaltail | -3,424 | 0,000 | 0,740 |
| RH_fimbria | 24,013 | 0,000 | 0,871 |
| RH_hippocampalfissure | 1,047 | 0,539 | 0,101 |
| RH_molecularlayerHP | 0,522 | 0,644 | 0,076 |
| RH_parasubiculum | -0,145 | 0,983 | 0,004 |
| RH_presubiculum | -2,000 | 0,000 | 0,592 |
| RH_subiculum | -0,457 | 0,180 | 0,217 |

#### Supplementary Figure 3

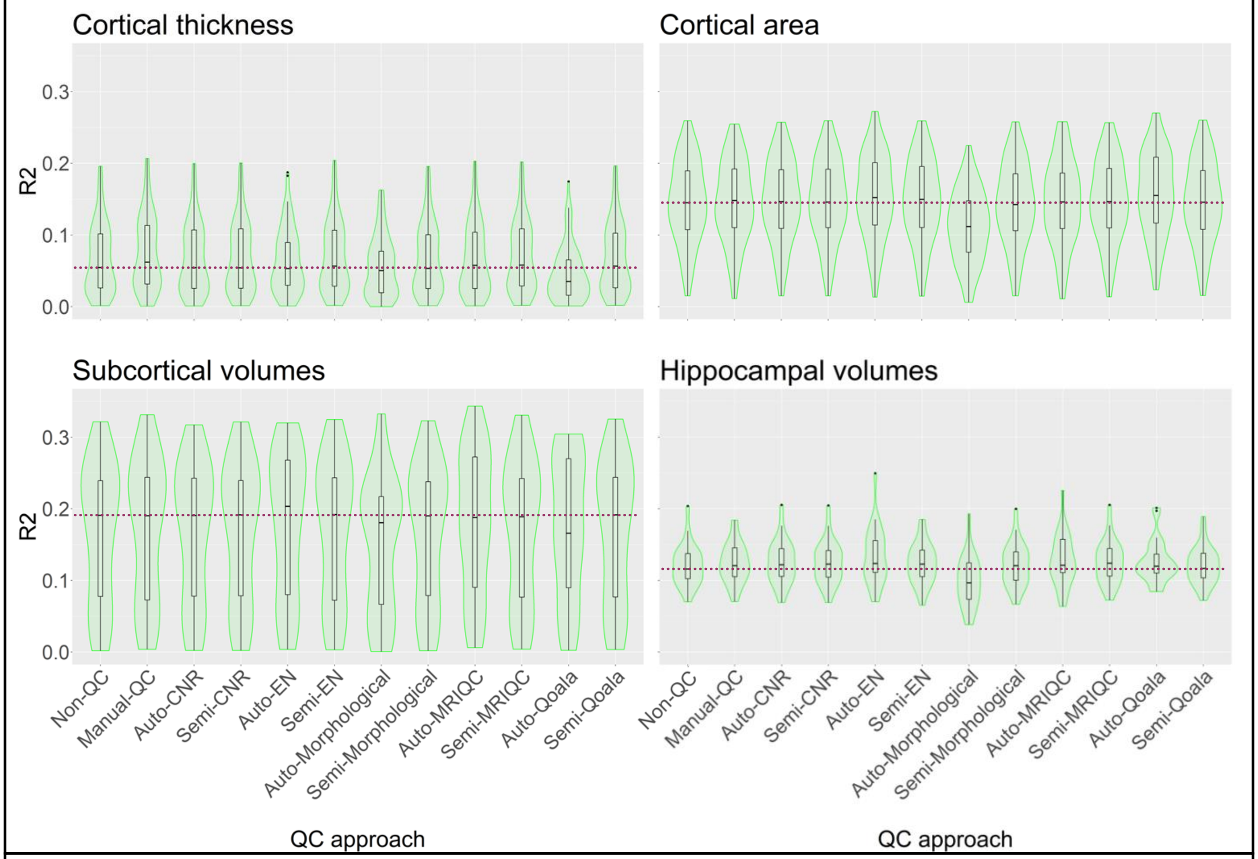


**Supplementary Figure 3:** Violin plots show the distribution of $R^2$ obtained by each QC strategy. The results are ordered by median and reported separately for each type of morphological measure: cortical thickness and cortical area (68 morphological measure each), subcortical volumes (38 morphological measures), and hippocampal subfields volume (24 morphological measures). Each $R^2$ point is the result of the linear model between age and a morphological measure, corrected for sex. Dotted line follows the median obtained by the Non-QC dataset.
